# Sugar availability suppresses the auxin-induced strigolactone pathway to promote bud outgrowth

**DOI:** 10.1101/752147

**Authors:** Jessica Bertheloot, François Barbier, Frédéric Boudon, Maria Dolores Perez-Garcia, Thomas Péron, Sylvie Citerne, Elizabeth Dun, Christine Beveridge, Christophe Godin, Soulaiman Sakr

## Abstract

- Apical dominance occurs when the growing shoot tip inhibits the outgrowth of axillary buds. Apically-derived auxin in the nodal stem indirectly inhibits bud outgrowth via cytokinins and strigolactones. Recently, sugar deprivation was found to contribute to this phenomenon.
- Using rose and pea, we investigated whether sugar availability interacts with auxin in bud outgrowth control, and the role of cytokinins and strigolactones, *in vitro* and *in planta*.
- We show that sucrose antagonizes auxin’s effect on bud outgrowth, in a dose-dependent and coupled manner. Sucrose also suppresses strigolactone-inhibition of outgrowth, and *rms3* strigolactone-perception mutant is less affected by reducing sucrose supply; however, sucrose does not interfere with the regulation of cytokinin levels by auxin, and stimulates outgrowth even with optimal cytokinin supply. These observations were assembled into a computational model where sucrose represses bud response to strigolactones, largely independently of cytokinin levels. It quantitatively captures our observed dose-dependent sucrose-hormones effects on bud outgrowth, and allows us to express outgrowth response to various combinations of auxin and sucrose levels as a simple quantitative law.
- This study places sugars in the bud outgrowth regulatory network, and paves the way for better understanding of branching plasticity in response to environmental and genotypic factors.

## INTRODUCTION

Shoot branching in plants is one of the major traits that affects the fitness of wild species in natural environments and the yield potential of agricultural, horticultural, and forestry crops (Evers *et al.*, 2011; Pierik & Testerink, 2014; Mathan *et al.*, 2016). Shoot branching markedly depends on the outgrowth of dormant or very slow growing axillary buds that form in the axils of leaves. Bud outgrowth is a highly plastic process, whose regulation involves a complex network of several interacting endogenous and exogenous cues (Rameau *et al.*, 2015).

Apical dominance is the term used to describe the inhibitory effect that the growing shoot tip exerts, at a distance, over the outgrowth of the axillary buds below. This systemic regulation is demonstrated by experiments where bud outgrowth is released after shoot tip decapitation.

Auxin, a plant hormone produced in the apical region and transported downwards through the stem, was considered important in the maintenance of apical dominance (Ongaro & Leyser, 2008). Indeed, exogenous auxin applied to the decapitated shoot tip can often restore bud outgrowth inhibition (Thimann & Skoog, 1933). However, auxin alone is insufficient to explain apical dominance. Firstly, for particular species and growing conditions, the supply of exogenous auxin to decapitated plants cannot completely restore apical dominance, suggesting that a factor other than auxin is involved (Cline, 1996). Secondly, correlative studies have shown that auxin transport, typically at 1 cm per hour through the stem, is too slow for local auxin depletion, following decapitation, to precede the onset of outgrowth of the basal bud in garden pea (Morris *et al.*, 2005; Renton *et al.*, 2012).

A recent study in pea (Mason *et al.*, 2014) indicated that the high demand for sugars by the growing shoot tip is an essential regulator of apical dominance. Following decapitation of the growing shoot tip, sugars rapidly redistributed (moving at ~150 cm per hour) and accumulated in the basal node and bud, prior to the onset of bud outgrowth, and while auxin levels in the adjacent node remained unchanged. This indicated that sugars might be the initial trigger of bud outgrowth after decapitation. This hypothesis was confirmed in the same study by showing that exogenous sugar supply through the petiole of plants with intact growing shoot tips was sufficient to induce bud outgrowth despite the presence of auxin in the stem. Furthermore, decreasing sugar levels through defoliation in decapitated plants delayed bud outgrowth. Additional studies in other species also support a role for sugars in apical dominance. Partial defoliation of sorghum plants, which reduced the number of sugar sources and lowered sugar levels in the bud, inhibited bud outgrowth (Kebrom & Mullet, 2015). The *tin* mutant of wheat, which has enhanced stem growth, and thus demand for sugars, shows reduced tillering (Kebrom *et al.*, 2012; Kebrom & Mullet, 2015). Sugars are proposed to play a signaling role in bud outgrowth regulation (Rabot *et al.*, 2012; Barbier *et al.*, 2015a; Barbier *et al.*, 2015b). This may be mediated, at least in part, by trehalose 6-phosphate, whose level indicates sucrose availability in plants (Figueroa & Lunn, 2016; Fichtner *et al.*, 2017).

While these data highlight that sugars and auxin are critically important mediators of apical dominance, until now, the roles of auxin and sugars in the regulation of bud outgrowth have been studied independently, and whether these two pathways interact during this process is still an open question.

Auxin in the main stem acts indirectly on lateral buds as it does not enter the bud (Booker *et al.*, 2003) and potentially act via two mechanisms involving cytokinins and strigolactones (Domagalska & Leyser, 2011 for review). In the first mechanism known as “the auxin canalization theory”, auxin in the stem acts via preventing the establishment and maintenance of auxin flow from axillary buds, a process promoting bud outgrowth (Prusinkiewicz *et al.*, 2009; Balla *et al.*, 2011). Within this, strigolactones and cytokinins respectively inhibit and promote auxin transport and export from axillary buds to the main stem (Shinohara *et al.*, 2013; Waldie & Leyser, 2018). However, recent findings in garden pea indicate that auxin canalization out of the bud is not involved in the initial stage of bud outgrowth, but that it would rather affect the sustained growth of already activated buds (Chabikwa *et al.*, 2018).

In the second mechanism, called “the second messenger theory”, auxin regulates the production of cytokinins and strigolactones that respectively induce or inhibit bud outgrowth (Sachs & Thimann, 1967; Gomez-Roldan *et al.*, 2008; Umehara *et al.*, 2008). Indeed, cytokinin biosynthesis and levels are rapidly enhanced in the nodal stem by auxin depletion (induced *e.g*. by decapitation, stem segment excision, application of an auxin transport inhibitor), and these phenomena can be prevented by exogenous auxin (Nordstrom *et al.*, 2004; Tanaka *et al.*, 2006). On the other hand, the expression of strigolactone biosynthesis-related genes is rapidly repressed by auxin depletion in the stem, a behavior that is also prevented by exogenous auxin application (Foo *et al.*, 2005; Zou *et al.*, 2006; Hayward *et al.*, 2009). Cytokinins and strigolactones are partly integrated within the bud by the transcription factor BRC1, involved in bud dormancy in several species (Aguilar-Martinez *et al.*, 2007; Dun *et al.*, 2012; Rameau *et al.*, 2015; Wang *et al.*, 2019).

Interestingly, sugars have been reported to have an opposite effect to auxin on cytokinins and strigolactones in different developmental processes (Arrom & Munne-Bosch, 2012; Li *et al.*, 2016; Tian *et al.*, 2018) including in bud outgrowth (Barbier *et al.*, 2015b; Barbier *et al.*, 2019). In our previous study on rose isolated nodal segments grown without auxin (Barbier *et al.*, 2015b), we highlighted that sucrose stimulated bud outgrowth, and that this growth was preceded by down-regulated strigolactone signaling gene expression and increased cytokinin synthesis. This is opposite to the effects of auxin on cytokinin and strigolactone biosynthesis (Tanaka *et al.*, 2006; Hayward *et al.*, 2009). These correlative trends indicate that increased sugar availability may antagonize auxin during the control of bud outgrowth and place strigolactones and cytokinins as potential integrators of such antagonism (Fig. 1). In this paper, we used physiological experiments to determine if and how sucrose, the main transported form of sugar in plants, and auxin interact to control bud outgrowth. Then, we tested the ability of this qualitative sugar-auxin interacting network to quantitatively reproduce the observed data by computer modelling, and derived from this model a simple law synthesizing the diversity of bud outgrowth response to the various combinations of sucrose and auxin levels.

**Fig. 1.**
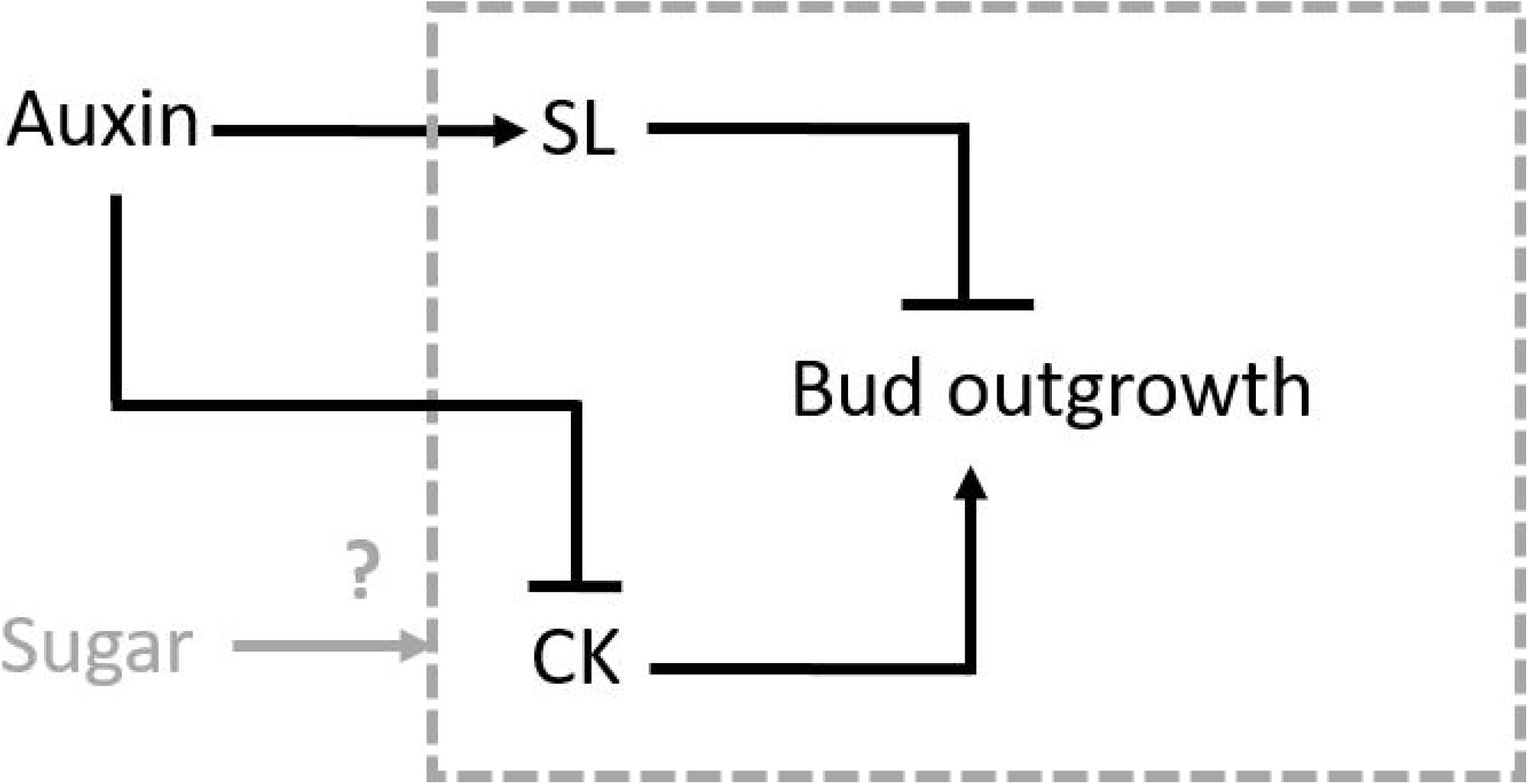
The interaction between sugar and auxin in the control of bud outgrowth is an open question. Auxin represses cytokinins (CK) and stimulates strigolactones (SL), which are stimulators and repressors of bud outgrowth, respectively (Beveridge *et al.*, 2000; Sorefan *et al.*, 2003; Nordstrom *et al.*, 2004; Foo *et al.*, 2005; Tanaka *et al.*, 2006; Zou *et al.*, 2006; Hayward *et al.*, 2009) (black). We test whether sugar interacts with auxin to control bud outgrowth via strigolactones and/or cytokinins (grey).

## MATERIALS AND METHODS

### Plant material and treatments

Rose plants were primary axes of *Rosa hybrida* L. cv. Radrazz obtained from cuttings. Pea plants were *Pisum sativum* L. cv. Terese (WT or *rms3* mutant) obtained from seeds. Environmental conditions for all experiments are described in supplemental table 1. Over-branched *rms3* mutants were grown under very low light intensity (70-80 μmol. m^−2^. s^−1^) to maintain buds in a state of dormancy until the transfer of nodal segments to *in vitro* conditions.

*In vitro* experiments involved the growth of nodal segments on MS medium supplemented with different concentrations of sucrose (10, 50, 100, 250 mM), 1-naphthaleneacetic acid (NAA; 0, 1, 2.5 μM), rac-GR24 (5 μM), and 6-benzylaminopurine (BAP; 10 μM), previously used in *in vitro* studies (Chiou & Bush, 1998; Rabot *et al.*, 2012; Barbier *et al.*, 2015b; Waldie & Leyser, 2018). For BAP, 10 μM is the optimal concentration that stimulates bud outgrowth for rose (Fig. S4). Except for Fig. **4b**, nodal segments were excised from the third true leaf-bearing node when the floral bud became visible for rose, and when the fourth true leaf was fully expanded for pea, and were placed on horizontal plates. For Fig. **4b**, nodal segments were excised from nodes five and six from plants with five or six expanded true leaves, and placed in upright open tubes. Details are given in supplemental text 1.

Experiments on decapitated plants of rose involved cutting 2 cm above the fourth leaf when the floral bud became visible. For all experiments, NAA (10 μM) was supplied in a basic medium to the decapitated stump. In Fig. **2a**, plants were either non-defoliated or partially defoliated. In Fig. **2b**, plants were (i) partially defoliated, and supplied at the second topmost leaf with mannitol (50 mM) or sucrose (50 mM), (ii) partially defoliated except at the second topmost leaf. For Fig. **4c**, plants were partially defoliated and vascularly supplied with GR24 (5 μM) 1 cm below the second downmost node (Fig. **4c**). Partial defoliation consisted of removing, four out of the five leaflets at each node. The methods for supply of hormones and sugars are described in supplemental text 2.

**Fig. 2.**
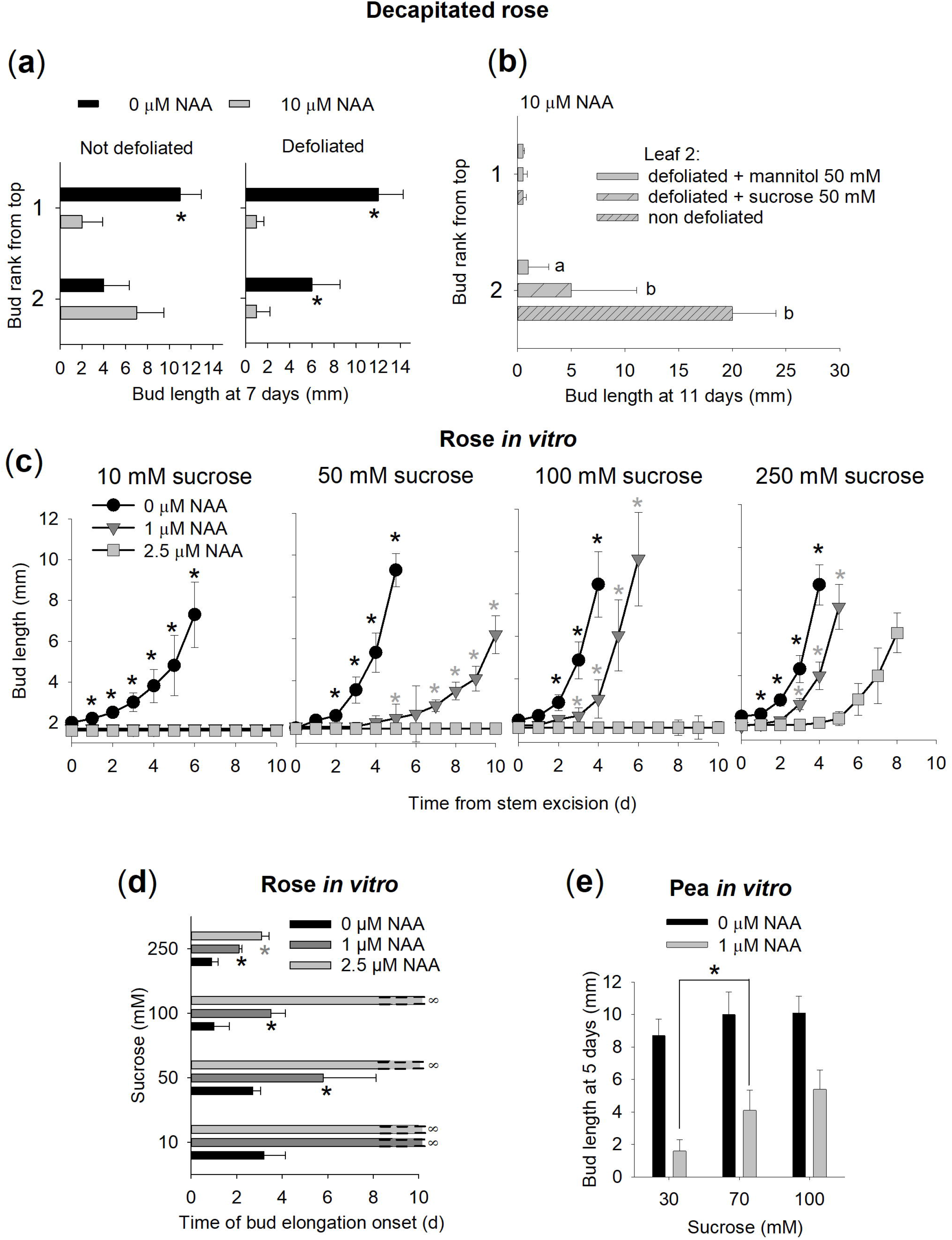
Sugars and auxin act antagonistically, and in a coupled and dose-dependent manner in the control of bud outgrowth. (**a**) Effect of NAA on the length of the two topmost buds of rose plants previously decapitated and either non-defoliated or partially defoliated. Data are medians (*n*=6). Asterisks indicate significant differences between NAA treatments (Wilcoxon’s test; *p*<0.05). (**b**) Bud length response to sucrose supply to the second topmost leaf of partially defoliated and decapitated plants supplied with NAA, compared to a mannitol supply (osmotic control), and compared to a plant where the second leaf is not defoliated. Only the two topmost buds are represented. Data are medians (*n*=8). Letters indicate significant differences between treatments (Wilcoxon’s test; *p*<0.05). (**c**, **d**) Bud outgrowth response to NAA for nodal segments of rose grown *in vitro* with increasing levels of sucrose: (c) elongation kinetics of the bud with the median final length (*n*=10); (d) median time at which elongation starts, unclosed horizontal bars (∞) representing no bud outgrowth. Black asterisks indicate significant differences between 0 and 1 μM NAA; grey asterisks indicate significant differences between 1 and 2.5 μM NAA (Wilcoxon’s test; *p*<0.05). (**e**) Bud outgrowth response to NAA for nodal segments of pea grown *in vitro* with increasing levels of sucrose. Data are medians (*n*=9). The asterisk indicates a significant difference between sucrose treatments (Wilcoxon’s test; *p*<0.01). For all graphs, error bars represent 95% confidence intervals.

### Bud outgrowth

Buds *in vitro* were photographed daily and bud length was quantified using ImageJ software (https://imagej.nih.gov/ij/). Rose buds display a phase of slow elongation followed by a phase of rapid elongation (Barbier *et al.*, 2015b). A bud was considered to grow out if it entered a phase of rapid elongation. The time at which the bud grew out was estimated as described in supplemental text 3. For decapitated rose plants, the state of each bud (outgrowing or not) along the stem and bud length was measured daily. A bud was considered to grow out if at least one visible leaf protruded between the two bud scales.

### Cytokinin concentrations

Cytokinin content of the nodal stem was determined as previously described (Barbier *et al.*, 2015b) (retention times, limits of quantification and detection in supplemental table 2). Nodal stem was defined as the bud and nodal segments with 5 mm of stem of each side of the node.

### Statistical analysis

Statistical analyses were done using R software for Windows. The functions t.test(), wilcox.test(), and fisher.test() were used for Student’s, Wilcoxon’s, and Fisher’s test, respectively.

### Model equations

The computational model is schematically described in Fig. **5a**, and model parameters are listed in supplemental table 3.

The levels of auxin (*A*) and sucrose (*S*) control the synthesis of cytokinins (*CK*) and strigolactones (*SL*), and the rates of change in the levels of cytokinins and strigolactones within a time step (*dt*) were described using a system of ordinary differential equations:

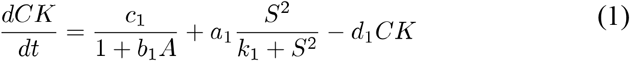

where *c*_1_, *b*_1_, *a*_1_, *k*_1_ and *d*_1_ are constants (see sup. table 3 for definition, units and values). The first term corresponds to auxin-repressed cytokinin synthesis, the second term sucrose-stimulated cytokinin synthesis (effects are supposed to be cumulative), and the last term cytokinin-dependent cytokinin degradation.

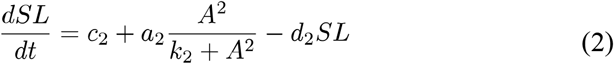

where *c*_2_, *a*_2_, *k*_2_ and *d*_2_ are constants (see sup. table 3 for definition, units and values). The first term is the base synthesis rate, the second term is auxin-stimulated synthesis, and the last term is strigolactone-dependent degradation.

Cytokinins and strigolactones control the synthesis of the signal integrator *I*, and *I* changes within a time step as follows:

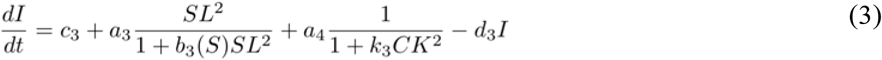

with

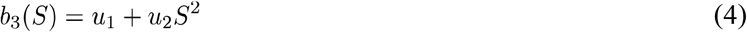

and where *c*_3_, *a*_3_, *a*_4_, *k*_3_, *d*_3_, *u*_1_, and *u*_2_ are constants (see sup. table 3 for definition, units and values). The first term is the base synthesis, the second term is strigolactone response, the third term is cytokinin response, and the last term *I*-dependent degradation. Inhibitor response is an increasing function of strigolactone level and is repressed by sucrose level. It is also a decreasing function of cytokinin level.

We assume in addition that the level of *I* correlates with the time at which bud outgrowth starts (*T*). A threshold (*I*_*0*_) determines if *T* is finite or infinite (bud outgrowth completely prevented), as follows:

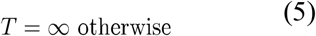

The model was implemented in Python (https://www.python.org/).

### Model calibration

Our model takes auxin and sucrose levels as an input and estimates corresponding delays in bud outgrowth. It relies on kinetic parameters whose values are in general not known. Here, we exploit our isolated bud system to indirectly estimate these kinetic parameters. For this, we experimentally fixed concentrations of input variables in the isolated bud medium, and measured values of output variables. We could then use these pairs of observed input/output observed values to estimate plausible values of the kinetic parameters. The imposed and measured values are detailed in supplemental table 4.

Based on different combinations of imposed input values, we used a gradient algorithm to infer model parameters. We varied the initial point in the parameter space by performing a sample of 1000 simulations starting from initial values of parameters randomly selected in a range of [0,1000] for all parameters except for decay parameters selected in a range of [0,1[. The gradient algorithm was achieved using the function least squares of the module scipy to estimate each parameter value (http://scipy.org/). The function optimized the different parameters of the model by minimizing the relative errors between measured values and estimated values of *CK* and *T*. Estimated parameter values are listed in supplemental table 3.

For each simulation, the algorithm converged to an optimized set of parameter values associated with a least square error threshold (0.60±1e-6). Interestingly, we observed that the optimized parameter values did not depend much on their initial value and had very close values (standard deviation <1e-2 for set of values of each parameter), suggesting that the numerical estimation of the parameters in this system is particularly robust.

## RESULTS

### Auxin and sugar availability control bud outgrowth in an antagonistic, coupled, and dose-dependent manner

We first evaluated the existence of an antagonistic effect of sugar supply and auxin in the regulation of axillary bud outgrowth. We performed physiological experiments using decapitated rose plants, a species in which we have previously established the action of sugars on bud outgrowth (Barbier *et al.*, 2015b), and manipulated levels of both auxin and sugar available to buds. Auxin levels were altered by treating the decapitated stump with or without NAA (1-naphthaleneacetic acid), a stable synthetic auxin. Sugar availability was manipulated by partial defoliation that is well-known to reduce plant sugar status (Kebrom & Mullet, 2015). The inhibitory effect of auxin was stronger in partially defoliated plants than in non-defoliated plants, in a manner that was negatively correlated with plant sugar status (Fig. **2a**, S1). While auxin only inhibited the topmost bud of non-defoliated plants, the second topmost bud was also inhibited in partially defoliated plants (Fig. **2a**), which have lower sugar levels than non-defoliated plants (Fig. S1). Defoliation could affect sugar status but also other physiological variables (*e.g.* transpiration stream, xylem-transported molecules; Cerasoll *et al.*, 2004; Lestienne *et al.*, 2006; Eyles *et al.*, 2013). Here, we show that sugar contributes to the bud outgrowth stimulation seen in non-defoliated plants compared to defoliated plants, because bud outgrowth was significantly induced at the second topmost node of defoliated plants when sucrose was supplied to its petiole or when its leaf was non-defoliated on the contrary to mannitol, an osmotic control (Fig. **2b**). This is in agreeance with the observation that auxin does not inhibit bud outgrowth in intact garden pea plants when sucrose is exogenously supplied (Mason *et al.*, 2014). These results support the idea that sugars and auxin regulate bud outgrowth in an antagonistic manner.

We then quantified the antagonistic effects of sugars and auxin on bud outgrowth using single nodal segments grown in split plates *in vitro*. This system has successfully been used in previous studies to easily manipulate the levels of several regulators of bud outgrowth (Chatfield *et al.*, 2000; Rabot *et al.*, 2012; Waldie & Leyser, 2018). The form of sugar used was sucrose, which is the main transported form of sugars in plants (Lemoine *et al.*, 2013). Sucrose concentration in the phloem sap greatly varies between plant species, ranging from 100-900 mM (Ohshima *et al.*, 1990; Nadwodnik & Lohaus, 2008; Jensen *et al.*, 2013). In peach, a rosacea species like rose, sucrose concentration in the phloem sap has been reported to be about 200 mM (Nadwodnik & Lohaus, 2008). The supply of 100 mM sucrose to rose nodal segments *in vitro* could antagonize the inhibiting effect of 1 μM NAA on bud outgrowth, which was not the case for 100 mM mannitol (Fig. S2).

In order to quantify the antagonistic effect of sucrose and auxin *in vitro*, we used sucrose concentrations ranging from 10 to 250 mM. Bud outgrowth is a continuous process, often measured as an elongation of the bud through time, which can be divided into a lag period before growth starts and a period of rapid growth (Chatfield *et al.*, 2000; Barbier *et al.*, 2015b). As done previously for rose, we quantitatively described bud outgrowth response by measuring whether or not buds grow out as well as the time at which their growth commenced (Barbier *et al.*, 2015b). In addition to rose, we here included garden pea, which has also previously been used to establish the importance of sugars in bud outgrowth (Mason *et al.*, 2014).

At low sucrose concentration (10 mM), rose buds grew out in the absence of NAA and were completely repressed by 1 and 2.5 µM NAA (Fig. **2c**). At intermediate sucrose concentrations (50 and 100 mM), high NAA (2.5 μM) completely suppressed bud outgrowth, while intermediate NAA (1 μM) only delayed the time at which elongation started (Fig. **2d**). This delay was inversely correlated to sucrose level (intermediate NAA delayed elongation by approximately 6 and 3.5 days for 50 mM and 100 mM, respectively). At a high sucrose concentration (250 mM), NAA was no longer able to completely suppress bud outgrowth (Fig. **2c**), but delayed the time at which elongation started in a dose-dependent manner (Fig. **2d**). This highlighted that sucrose only partially removed the inhibitory effect of auxin. This effect of sucrose was also observed for nodal segments of garden pea *in vitro* (Fig. **2e**).

Interestingly, the amplitude of the sucrose effect depended on auxin level. At high auxin (2.5 μM NAA) the effect of a given change in sucrose level on bud outgrowth was high (reduction in the time of elongation onset of more than 5% between 100 and 250 mM sucrose), while it remained intermediate (reduction of 2% between 50 and 250 mM sucrose) at intermediate auxin levels (1 μM NAA), and low (reduction of 1% between 10 and 250 mM sucrose) at 0 μM NAA. This shows that sucrose and auxin have a coupled effect on bud outgrowth. A two-way ANOVA analysis also indicated a significant interaction between auxin and sucrose on the time at which elongation starts (*p-value* < 10^−4^).

All these results indicate that sugar partially antagonizes the effect of auxin on bud outgrowth in a coupled manner, and that combined sugar and auxin levels quantitatively modulate bud outgrowth by determining whether buds grow and the time at which their growth starts. Current knowledge has led to a model where auxin in the nodal stem inhibits the early stage of bud outgrowth through modulation of cytokinin and strigolactone levels (Domagalska & Leyser, 2011). Since sucrose supply to rose nodal stem segments induces rapid changes in cytokinin levels and strigolactone signaling (Barbier *et al.*, 2015b), we sought to determine the role of these two hormones in the modulation of bud outgrowth by sugar-auxin interactions.

### Sugar availability modulates bud outgrowth independently of cytokinin levels

The suppression of nodal cytokinin content by auxin was the first hormonal mechanism proposed to explain the indirect action of auxin in apical dominance (Shimizu-Sato *et al.*, 2009). Our previous study demonstrated that sucrose supply to nodal segments of rose *in vitro* could upregulate cytokinin synthesis (Barbier *et al.*, 2015b). We therefore tested whether auxin and sucrose might antagonistically affect cytokinin levels in our isolated rose bud system. There was a substantial and widespread suppressive effect of NAA on endogenous cytokinins, regardless of the sucrose concentration in the growth medium (Fig. **3a**, Fig. S3). In the presence of 2.5 μM NAA, increasing sucrose from 100 to 250 mM caused only a small increase in the level of the intermediate forms of cytokinins, and no significant increase in the active form iP (Fig. **3a**), while inducing a clear positive response in bud outgrowth (Fig. **2c,d**). This contrast in impact of sucrose on bud outgrowth versus cytokinin levels was also observed in the presence of 1 μM NAA. In this case, increasing sucrose from 50 to 100 mM did not significantly increase cytokinin, while reducing the delay before bud elongation (Fig. **2c,d***vs.* Fig. S3). These results indicate that only a minor component of the stimulatory effect of sugar on bud outgrowth may occur via sugar modulation of cytokinin levels in the rose single node.

**Fig. 3.**
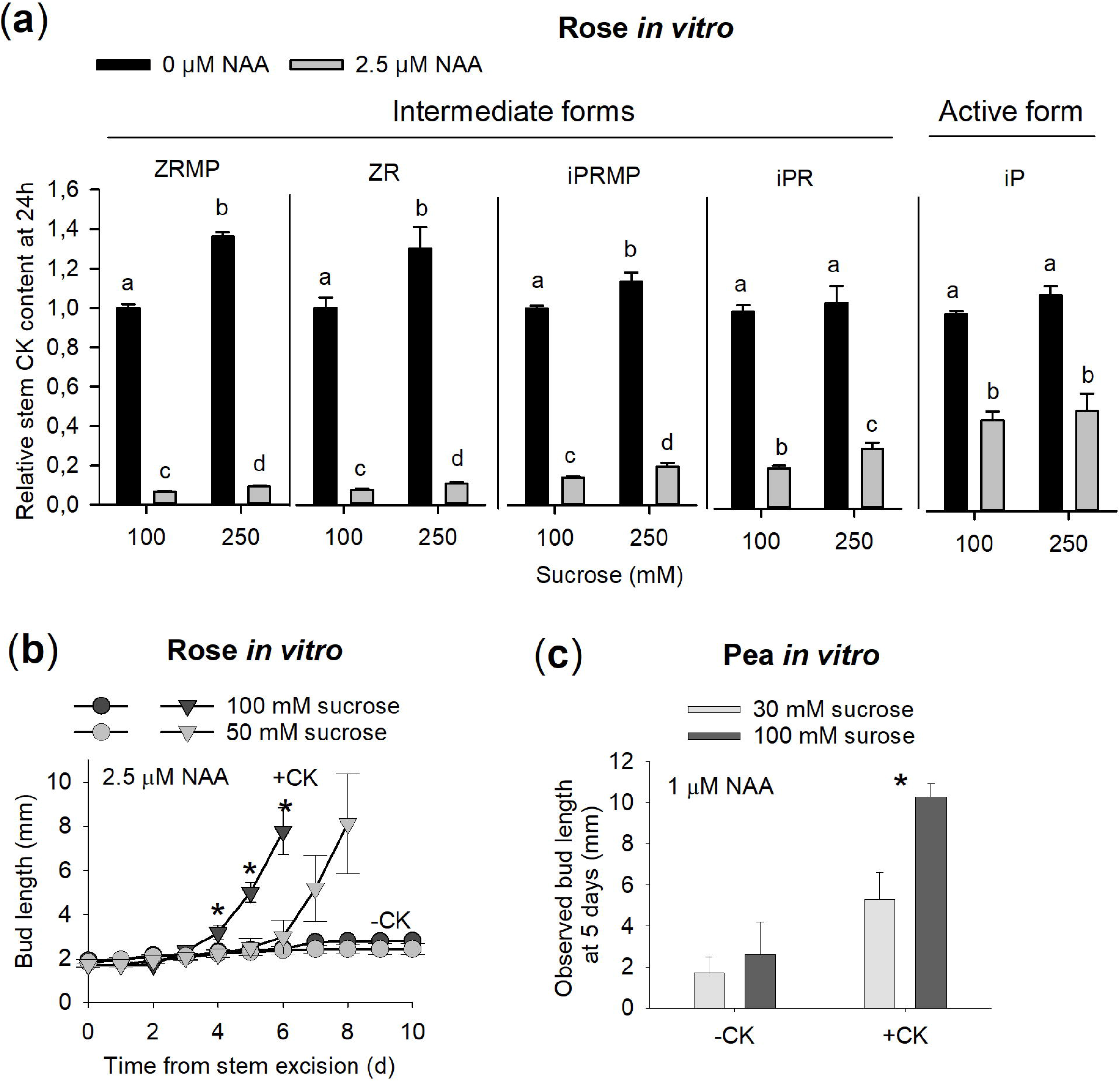
Sucrose acts independently of cytokinin levels. (**a**) Response of nodal cytokinins to NAA for nodal segments of rose grown *in vitro* with increasing levels of sucrose. Data are means ± SEM (*n*=4 pools of 3 stem segments). Expression data were measured 24h after nodal stem excision. Values are represented relative to the treatment 100 mM sucrose and 0 μM NAA. Different letters indicate significant difference between means (Student’s test; *p*<0.05). (**b**, **c**) Impact of BAP, a synthetic cytokinin (−CK/+CK), on the inhibition of bud outgrowth by NAA for nodal segments of rose (b) and pea (c) grown *in vitro* with two sucrose concentrations: (b) elongation kinetics of the bud with the median final length (*n*=10); (c) median bud length at 5 days (*n*=9). Error bars represent 95% confidence intervals. Asterisks indicate significant differences between sucrose treatments in presence of BAP (Wilcoxon’s test; *p*<0.01).

To confirm this result, we supplied NAA-inhibited buds with a concentration of synthetic cytokinin (6-benzylaminopurine -BAP- at 10 μM) that is optimal for bud outgrowth (above this concentration, there is no further stimulation of bud outgrowth; Fig. S4) and tested the impact of two sucrose concentrations on bud response. As expected, in the absence of cytokinins, 2.5 µM NAA inhibited buds under both 50 and 100 mM sucrose (Fig. **3b**). Cytokinin supply triggered bud outgrowth under both sucrose conditions but, interestingly, the time at which outgrowth started was sucrose-dependent (Fig. **3b**). Cytokinin treated buds elongated earlier under the higher sucrose concentration. Similarly for pea, cytokinin supply released buds from NAA inhibition at both 30 and 100 mM sucrose, and cytokinin-treated buds were longer at high than at low sucrose level (Fig. **3c**). Thus, even in the presence of exogenously supplied cytokinin, sucrose is still able to promote bud outgrowth. Combined with the observation that cytokinin levels only show a minor response to sucrose in presence of auxin, these data support the premise that sugar acts largely independently of cytokinin levels to stimulate bud outgrowth in presence of auxin.

### The sugar pathway acts by suppressing bud response to strigolactones

Previous studies have shown that sucrose does not repress the expression of strigolactone synthesis genes, but down-regulates the expression of a strigolactone signaling gene (Kebrom *et al.*, 2010; Barbier *et al.*, 2015b). We reasoned that sucrose may down-regulate or suppress the response of the bud to strigolactones, rather than regulating synthesis of strigolactones. To examine the effect of sugars in the strigolactone bud-inhibition response, we exposed rose nodal segments *in vitro* to different sucrose concentrations with an intermediate auxin concentration, that would potentially enable a strigolactone inhibition response (Crawford *et al.*, 2010). At 50 mM sucrose, the supply of GR24, a synthetic strigolactone, in the growth medium was able to greatly suppress bud outgrowth (Fig **4a**). However, at 100 mM sucrose, this effect was completely supressed. This shows that sucrose suppresses the strigolactone-inhibition of bud outgrowth.

**Fig. 4.**
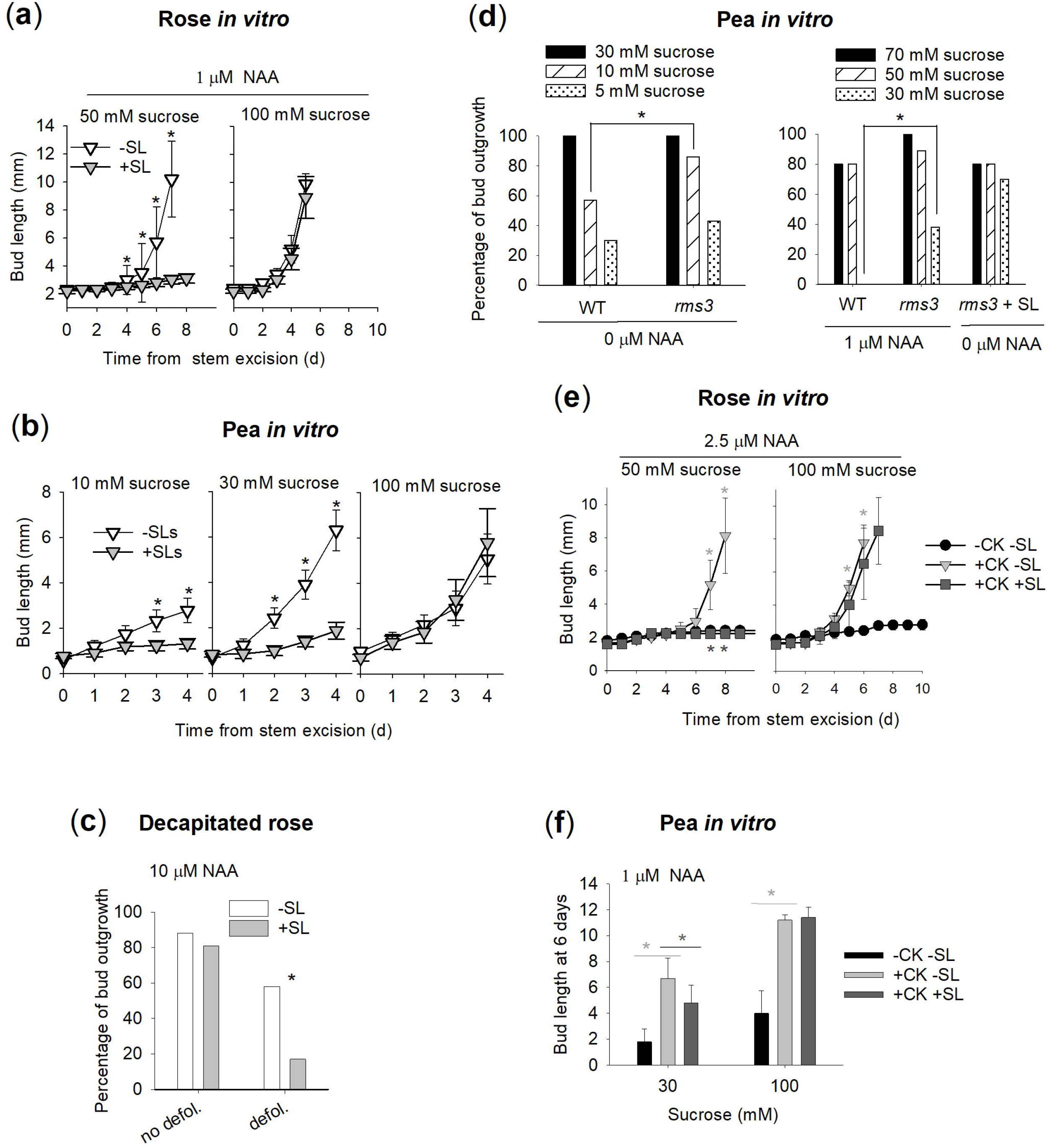
Sucrose suppresses bud outgrowth response to strigolactones, and *rms3* mutant displays a reduced response to a decrease in sucrose level. (**a**, **b**) Bud outgrowth response to GR24, a synthetic strigolactone (−SL/+SL), for nodal segments of rose (a) and pea (b) grown *in vitro* with increasing sucrose concentrations. Data represent the bud with the median final length (*n*=10 buds for rose and *n*=9 buds for pea). Asterisks indicate significant differences between GR24 treatments (Wilcoxon’s test; *p*<0.01). For rose, the medium is supplemented with NAA (1 μM). (**c**) Impact of GR24 supply (−SL/+SL) on bud outgrowth percentage of rose plants, previously decapitated and either non-defoliated (no defol.) or partially defoliated (defol.) (*n*=8). Only the buds above the point of GR24 supply are considered. Asterisks indicate a significant effect of GR24 (Fisher’s exact test; *p*<0.1). (**d**) Sucrose impact on the percentage of bud outgrowth for wild-type plants of garden pea, and *rms3* mutant deficient in a strigolactone signalling gene. Nodal segments were grown *in vitro* without or with NAA or GR24 (+SL), and a range of sucrose concentrations (5, 10, 30 mM under 0 μM NAA; 30, 50, 70 mM under 1 μM NAA). A bud was considered to grow out when its length was above 3 mm. Asterisks indicate significant differences between treatments (*n*=9 to 12; Fisher’s exact test; *p*<0.1). (**e, f**) Impact of supplying BAP (+CK −SL) and BAP plus GR24 (+CK +SL) on bud outgrowth for nodal segments of rose (e) and pea (f) grown *in vitro* at two sucrose concentrations: (e) observed elongation kinetics of the bud with the median final length (*n*=10); (f) observed median bud length at 6 days (*n*=9). Light grey asterisks indicate a significant effect of BAP supply (−CK −SL *vs.* +CK −SL), dark grey asterisks indicate a significant effect of GR24 supply in presence of BAP (+CK −SL *vs.* +CK +SL) (Wilcoxon’s test; *p*<0.05). For all graphs, error bars represent 95% confidence intervals.

Similar results were observed for pea, except in this case the addition of auxin in the media was unnecessary (Brewer *et al.*, 2015). GR24 had a significant inhibitory effect at 10 and 30 mM sucrose in pea, but was ineffective at 100 mM sucrose (Fig. **4b**), indicating that the ability of sugar to repress the bud response to strigolactones is conserved in diverse species.

To test this hypothesis *in planta*, we decapitated rose plants with different levels of leaf-supplied sugars modulated by defoliation, as done previously (Fig. **2c**, S1). GR24 was more effective at inhibiting bud outgrowth at high defoliation than without defoliation (Fig. **4c**).

To further test whether sugar inhibits strigolactone response to stimulate bud outgrowth, we compared the response of WT and the *rms3* strigolactone perception mutant (de Saint Germain *et al.*, 2016) to variations in sucrose concentrations, with or without NAA. These concentrations allowed us to have a gradient of bud outgrowth percentage for the WT (Fig. **4d**). Compared to WT, *rms3* bud outgrowth was less responsive to a decrease in sucrose concentration. At the highest sucrose concentration (30 and 70 mM for 0 and 1 μM NAA, respectively), WT and *rms3* exhibited 100% bud outgrowth in the presence and absence of NAA; however, in contrast to WT, bud outgrowth of *rms3* was not as inhibited when sucrose was reduced to 10 mM in the absence of NAA, or to 30 mM in the presence of NAA, or to 30 mM in the presence of GR24. This reduced response to a decrease in sucrose concentration in the strigolactone perception mutant *rms3* supports an involvement of strigolactone pathway in sugar-stimulated bud outgrowth.

Auxin inhibition of bud outgrowth involves an antagonistic effect between strigolactones and cytokinins (Domagalska & Leyser, 2011; Dun *et al.*, 2012). To determine if sugar disrupts the antagonistic action of strigolactones and cytokinin on bud outgrowth, we supplied BAP and GR24 to rose and pea nodal segments *in vitro* at two sucrose levels (50 or 30 mM for rose and pea, respectively, and 100 mM for both species) and observed bud outgrowth that ensued. NAA was supplied in a quantity sufficient to inhibit bud outgrowth in the absence of cytokinin for both species. As described previously (Fig. **3b**), adding BAP stimulated bud outgrowth at both sucrose levels (Fig. **4e,f**). However, adding GR24 antagonized the positive effect of BAP only at the lower sucrose level, and not at the higher sucrose level. This suggests that the pathway of strigolactones involved in auxin effect is not able to inhibit bud outgrowth in high sugar environments.

### A computational model, in which sugar suppresses strigolactone pathway, captures the diversity of dose-dependent observations in a quantitative manner

Taken together, our biological results indicate that the antagonism of sugar to auxin on bud outgrowth involves sugar suppression of strigolactone response. To check whether this hypothesis is quantitatively sufficient to explain the diversity of biological effects of sucrose and hormones on bud outgrowth, we constructed a computational model of our putative sugar-hormone network (Fig. **5a**) and tested its ability to quantitatively reproduce the range of phenotypes resulting from sucrose and hormone crosstalk experiments. The inputs of the model correspond to the levels of sucrose and auxin; the output is the time at which bud outgrowth starts, which is either infinite (no bud outgrowth) or initiates at different time points. The model relies on the following assumptions. According to literature, auxin (i) suppresses cytokinin synthesis (Tanaka *et al.*, 2006; Fig. **3a**, S2) and (ii) enhances strigolactone synthesis (Hayward *et al.*, 2009; Fig. S4). Cytokinins and strigolactones induce responses that are antagonistically integrated at the bud and control bud outgrowth (Dun *et al.*, 2012). According to our results, sucrose suppresses the strigolactone response (Fig. **4**) without significantly altering strigolactone synthesis (Barbier *et al.*, 2015b). On the other hand, sucrose causes only a small enhancement of cytokinin content (Fig. **3a**, S2). We modelled these interactions using a set of coupled ordinary differential equations to account for the quantitative variations of the different variables (see material and methods).

**Fig. 5.**
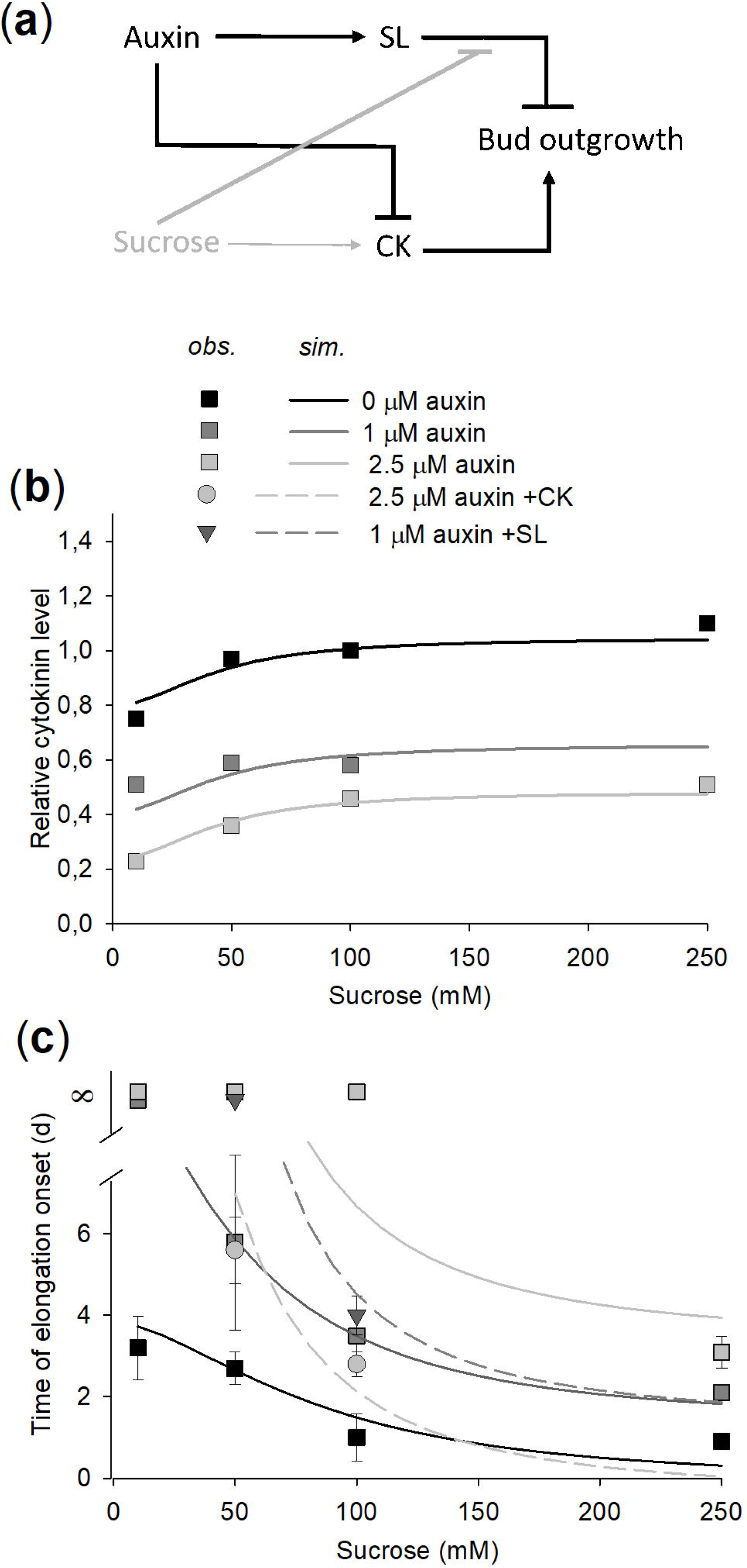
A model including sucrose as a suppressor of strigolactone response captures all observed crosstalks between sucrose and hormones. (**a**) A model of sucrose crosstalk with auxin in bud outgrowth control. Sucrose suppresses auxin-induced strigolactone pathway, and causes only a small increase in the level of auxin-repressed cytokinins. (**b**, **c**) Simulated (lines) and observed (symbols) nodal cytokinin levels (**b**) and times at which bud elongation starts (**c**) for nodal segments of rose grown *in vitro* at different sucrose levels, auxin levels, without or with cytokinins and strigolactones (+CKs, +SLs). In (c), observations of an absence of bud elongation are represented by an infinite time at which elongation starts (∞). For simulations, bud elongation is completely prevented above a threshold of 8.3 days. Observed cytokinin levels were calculated as detailed in supplemental table 4. Observed times at which elongation starts are those of Fig. **2d** and those calculated from Fig. **3b**, **4a** (see supplemental text 3 for calculation details).

As the kinetic parameter values involved in these equations are mostly unknown in the literature, we sought to estimate these values using our rose *in vitro* experiments. These experiments provided measurable outputs corresponding to controlled input levels. We then used these observed pairs of input/output to find the most plausible parameter values of the model to account for all our biological observations, namely the bud outgrowth responses to the different concentrations and combinations of sucrose, auxin, cytokinins, and strigolactones (Fig. **2**–**4**). For this, we used a systematic exploration of the parameter space, constraining the model to the observed endogenous cytokinin levels and to the observed time at which elongation starts for the available experimental data and treatments (supplemental text 6). From this analysis, we discovered a relatively narrow region of the parameter space where the model can optimally reproduce the observed interactions between sucrose and hormones. In this region, the model captured the conditions of hormone and sucrose levels for bud elongation as well as the time at which bud elongation starts (Fig. **5c**). In particular, it accounted for the sucrose x auxin interaction effect that was observed in the time at which outgrowth starts.

### Bud outgrowth is controlled by a simple variable combining both sucrose and auxin levels

Our modelling and experimental work shows that different combinations of sucrose and auxin levels can result in identical (or close to identical) bud outgrowth responses (*e.g.*, similar outgrowth response time for 1 μM NAA/100 mM sucrose and for 2.5 μM NAA/250 mM sucrose). This results from the antagonistic effect of the two input factors: for example, starting from given levels of auxin and sucrose, increasing the auxin level (*i.e*. increasing inhibition) can be compensated for by an adequate increase in the sucrose level (increasing bud outgrowth release). We wondered whether we could extract a law from the model to help us quantitatively predict how to maintain balance between the two antagonistic factors. Based on the model’s equations at equilibrium, we analyzed different algebraic combinations of the input variables, and found one that made it possible to summarize the overall system’s behaviour using a simple combination of the input levels of auxin (*A*) and sucrose (*S*):

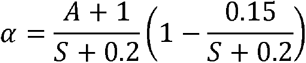

This variable, *α*, combines auxin and sucrose levels so that each value of *α* defines a unique time at which bud outgrowth starts, through a close to linear function of *α* (Fig. **6**). Interestingly, other combinations of auxin and sucrose levels would not lead to a similar one to one relationship (Fig. S4). We call *α* a control variable for bud outgrowth. This control variable allows us to summarize efficiently the behaviour of the system without needing to know or to run the model: at high sucrose levels, the time at which outgrowth starts is basically a linear function of the ratio between simple affine functions of auxin and sucrose levels 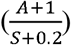; at low sucrose levels, this ratio is decreased by a correcting term 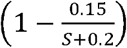.

**Fig. 6.**
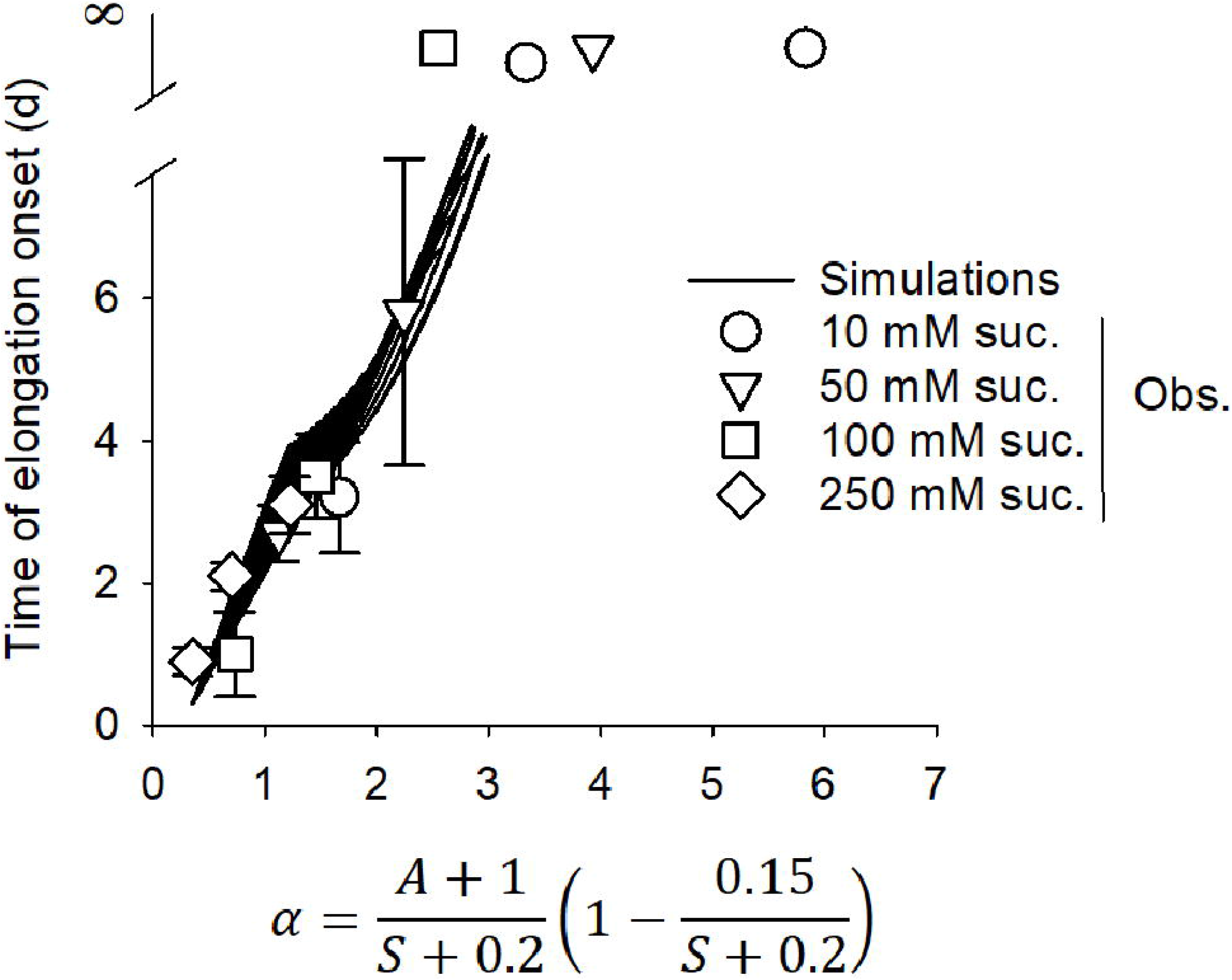
Bud outgrowth is controlled by a simple variable combining both sucrose and auxin levels. Relationship between (i) the time at which elongation starts and (ii) the ratio between affine functions of auxin and sucrose levels 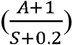, modulated by a sucrose-dependent correcting term 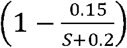. The different lines show simulations for different sucrose levels and the different symbols the biological observations for nodal segments of rose *in vitro* grown at different sucrose levels (Obs.). Observations of an absence of bud elongation are represented by a time at which elongation starts which is infinite (∞). For simulations, bud elongation is completely prevented above a threshold of 8.3 days.

## DISCUSSION

Apical dominance results in growth in height at the expense of lateral growth by inhibiting axillary buds. In the classical view, auxin is a signal that indicates the presence of growing apical organs and that inhibits the outgrowth of axillary buds at nodes below (Ongaro & Leyser, 2008). Together with results demonstrating sugars as a positive signal for bud outgrowth (Rabot *et al.*, 2012; Barbier *et al.*, 2015b; Fichtner *et al.*, 2017), the recent study by Mason et al. (Mason *et al.*, 2014) on garden pea has drawn attention to the role of the growing apical shoot tip in creating a sugar demand which diverts sugars from the axillary buds and inhibits their outgrowth. Using model species rose and pea, and isolated nodal segments and decapitated plants to modulate both sugar and auxin levels for buds, we show that bud outgrowth is under an antagonistic coupled control of sugar and auxin levels. More precisely, the ratio between simple functions of sugar and auxin levels determines if a bud grows out and the time at which growth starts (modulo a correcting term at low sugar level). These results bring the view that plant sugar status modulates auxin-related apical dominance. In this view, auxin produced by the growing shoot tip results in strong apical dominance only if sugar status of the plant is low. When plant sugar status is high, apical dominance is reduced, leading to bushy phenotypes. Consistently, a recent transcriptomic study highlighted that bud dormancy is, at least partly, maintained by a carbon starvation syndrome in annual and perennial plants (Tarancon *et al.*, 2017). Moreover, bud outgrowth is influenced by modulating either sources (exogenous sugar supply, partial defoliation, CO_2_ supply) or sinks (decapitation, *tin* mutation) for sugars (Kebrom *et al.*, 2012; Mason *et al.*, 2014; Kebrom & Mullet, 2015; Fichtner *et al.*, 2017; Kebrom, 2017; Otori *et al.*, 2017; Martin-Fontecha *et al.*, 2018).

Sugars could take part in the strong response of plant branching to a myriad of environmental conditions (Rameau *et al.*, 2015). Indeed, environment impacts the source-sink balance within the plant (Kebrom, 2017) and thus its sugar status. Under environmental conditions leading to high source-sink balance, plant sugar status is increased and this could result in reduced auxin-related apical dominance and induction of bud outgrowth. In support, experiments have shown that the efficacy of exogenous auxin to inhibit bud outgrowth was reduced in growing conditions such as high light intensity, which promotes photosynthesis and sugar status (Gregory & Veale, 1957; Cline, 1996). Sugar status would then provide an internal cue to enable an optimized response to the environment, coordinating investment in lateral growth relative to the whole plant. The dose-dependent effect of sugars that we highlight here could provide a means for plants to fine-tune their architecture.

Surprisingly, we found little evidence that cytokinin levels mediate the antagonistic effect of sugar to auxin. Cytokinins are positive endogenous signals responsible for the stimulation of tissue sink strength (Roitsch & Ehness, 2000; Werner *et al.*, 2008; Roman *et al.*, 2016a) and of bud outgrowth by light or nitrogen nutrition (Takei *et al.*, 2002; Kamada-Nobusada *et al.*, 2013; Roman *et al.*, 2016b; Corot *et al.*, 2017). Sugars stimulate cytokinin biosynthesis and/or level in different physiological processes (Barbier *et al.*, 2015b for review). However, we report here that sugar does not antagonize the strong repressing effect of auxin on cytokinin levels at concentrations where it reduces bud inhibition by auxin. In addition, sugar is still able to promote bud outgrowth in presence of exogenously supplied cytokinins. Such non-involvement of cytokinins in the antagonistic effect of sugar to auxin on bud outgrowth is consistent with the result of a recent experiment using cytokinin deficient mutants of Arabidopsis (Muller *et al.*, 2015). In this experiment, decapitation, which is thought to stimulate bud outgrowth by increasing plant sugar status, led to highly-branched phenotypes for both wild-type plants and cytokinin deficient mutants. Additionally, excised single nodes of these mutants did not display any increased responsiveness to auxin when grown *in vitro* on medium containing sucrose (Muller *et al.*, 2015). Consequently, modulation of cytokinin levels was clearly not critical for the decapitation response and for sucrose-dependent bud outgrowth (Barbier *et al.*, 2019). However, further study should clarify whether sugar could affect cytokinin signalling to regulate bud outgrowth, as it does in regulation of root growth (Kushwah *et al.*, 2011; Kushwah & Laxmi, 2017).

We highlight that sugar supply inhibits strigolactone response to promote bud outgrowth. Strigolactones inhibit bud outgrowth and mediate the effect of auxin (Beveridge *et al.*, 2000; Zou *et al.*, 2006; Hayward *et al.*, 2009) as well as the response to different abiotic stresses which modulate strigolactone synthesis (phosphate or nitrogen deficiency) or signaling (drought) (Umehara *et al.*, 2010; Kohlen *et al.*, 2011; Bu *et al.*, 2014; Ha *et al.*, 2014; Saeed *et al.*, 2017). We show that sugar supply is able to repress the inhibitory effect of strigolactones on buds, as it is the case for strigolactone-induced bamboo leaf senescence in dark (Tian *et al.*, 2018). Moreover, *rms3*, a strigolactone-perception pea mutant, exhibits a reduced inhibition with decreasing sucrose concentration. This holds also true for seedling development of *max2,* a strigolactone signaling mutant, which displayed a reduced response to sugar (Li *et al.*, 2016). This effect of sugar on bud outgrowth through strigolactone pathway matches with the sucrose-mediated repression of *MAX2* expression in rose buds (Barbier *et al.*, 2015b) and the downregulation of *MAX2* in response to defoliation and shade in sorghum (Kebrom *et al.*, 2010). Moreover, in rose and pea, sucrose inhibited the expression of *BRC1* (Mason *et al.*, 2014; Barbier *et al.*, 2015b), encoding a transcription factor inhibiting bud outgrowth (Aguilar-Martinez *et al.*, 2007) and also involved in strigolactone signaling (Braun *et al.*, 2012; Dun *et al.*, 2012; Dun *et al.*, 2013; Seale *et al.*, 2017). Collectively, these findings prompt us to identify the molecular components of strigolactone signaling involved in sugar-mediated bud outgrowth promotion.

Previous results highlighted an effect of sugar on auxin signaling pathways on bud outgrowth (Rabot *et al.*, 2012; Barbier *et al.*, 2015b; Fichtner *et al.*, 2017). We further propose a simple model in which sugar and auxin interact in bud outgrowth regulation through modulation of the balance between cytokinins and strigolactones; this balance is a quantitative regulator that determines whether a bud grows out and the time at which it grows out. This simple model is sufficient to capture the variety of bud outgrowth responses *in vitro* to sucrose level, auxin level, and cytokinins and strigolactones. Like all models, it is a simplification of the physiological reality and it does not exclude the involvement of other mechanisms. In particular, it does not explicitly account for the role of auxin canalization out of the bud in controlling its outgrowth. However, auxin canalization is not involved in early outgrowth regulation (Chabikwa *et al.*, 2018), and could be considered as a mechanism downstream of cytokinins and strigolactones signaling such as BRC1 (Dun *et al.*, 2012), since both hormones also regulate canalization (Shinohara *et al.*, 2013; Waldie & Leyser, 2018).

In conclusion, we demonstrate that bud outgrowth quantitatively adjusts to the balance between sugar and auxin level, with increased sugar leading to a strong reduction of bud inhibition by auxin, and that the sugar effect involves repression of strigolactone response. As mentioned above, high sugar levels may explain a reduction of apical dominance in response to environmental or genetic factors increasing the source/sink balance within the plant. Sugars act through different signaling pathways, including those related to sucrose-, hexokinase-, glycolysis (Li & Sheen, 2016; Sakr *et al.*, 2018; Wingler, 2018). It has been suggested that sucrose could promote bud outgrowth in a signaling manner in rose and pea (Barbier *et al.*, 2015b; Fichtner *et al.*, 2017) and further experiments are required to determine the molecular mechanisms underlying the interactions between sugar availability and auxin. In addition to sugar, cytokinins and strigolactones have been shown to be involved in the response of branching to several environmental factors (Takei *et al.*, 2002; Drummond *et al.*, 2015; Roman *et al.*, 2016b; Saeed *et al.*, 2017). We suggest that our network involving auxin, sugars, cytokinins, and strigolactones may be a key integrator of the plant growth status and environmental conditions, to dynamically adapt plant architecture and thus contribute to plant plasticity.

## ACKNOWLEDGMENTS

We thank Bénédicte Dubuc and ImHorPhen team (UMR IRHS, France) for rose production, Hervé Autret for computer help, Catherine Rameau and François-Didier Boyer (IJPB, Versailles, France) for providing the seeds of *Pisum sativum* and GR24. We thank the Environment and Agronomy department of INRA for financial support.

## AUTHOR CONTRIBUTIONS

J. Bertheloot, C. Godin, and S. Sakr designed the research.

J. Bertheloot, F. Barbier, MD Perez-Garcia, T. Péron, S. Citerne, performed the experiments and analysed the data.

J. Bertheloot, F. Boudon, and C. Godin designed the model and the link between experiments and modelling, and analysed the simulations.

J. Bertheloot, F. Barbier, F. Boudon, E. Dun, C. Beveridge, C. Godin, S. Sakr contributed to manuscript writing.

J. Bertheloot and F. Barbier contributed equally to the work.

## SUPPORTING INFORMATION

**Supplemental text 1.** *In vitro* cultivation

**Supplemental text 2.** Exogenous supply of hormones in rose decapitated plants.

**Supplemental text 3.** Estimation of the time at which growth starts.

**Supplemental text 4.** Sugar content determination.

**Supplemented table 1.** Growth environment for each experiment.

**Supplemental table 2.** Cytokinin quantification: retention times, limit of quantification (LOQ), and limit of detection (LOD)

**Supplemented table 3.** Definition, units, and estimated values of the model parameters.

**Supplemental table 4.** Values of the variables used for model calibration.

**Fig. S1.** The level of defoliation of decapitated plants modulates sugar level.

**Fig. S2.** Sucrose antagonizes the inhibiting effect of auxin on bud outgrowth for rose nodal stems *in vitro*.

**Fig. S3.** Cytokinin levels of the nodal stem strongly change with auxin, but not with sucrose.

**Fig. S4.** 10 μM BAP optimally stimulates bud outgrowth for rose nodal segments *in vitro*.

**Fig. S5.** Deviations from the one-to-one relationship after changes of the control variable *α*.

