## Supplemental Material for "Sugar availability suppresses the auxin-induced strigolactone pathway to promote bud outgrowth"

**Supplemental text**

**Supplemental text 1. *In vitro* cultivation**

For all experiments except that of Fig. **4b**, 7 mm of stem on either side of the node were excised from plants. The leaf was removed, and cuttings were then disinfected in ethanol (ethanol/water; v/v; 70/30) for 10 to 20 s and transferred to horizontal plates (12 cm × 12 cm square Petri dishes) under a laminar flow hood. Plates had been previously filled with an autoclaved basic culture medium [full strength Murashige and Skoog medium, 1% agar, and 2.5 ml l^-1^ Plant Preservative Mixture (PPM; Kalys)] supplemented with different conditions of sucrose and hormones (NAA, BAP, rac-GR24). A 10 mm-wide strip of the medium had been removed from the plates to dig a gap. Nodal segments were placed so that both ends of the node cutting touched the medium.

For Fig. **4b**, 10 mm of stem on either side of the node were excised from plants. The leaf was removed. Each end of the nodal segment was embedded into an open tube (Brewer *et al.*, 2015) containing a basic culture medium (full strength Murashige and Skoog medium, 0.9% agar, and 0.1% PPM) supplemented with sucrose and rac-GR24. Nodal segments were placed upright.

**Supplemental text 2. Exogenous supply of hormones in rose decapitated plants.**

NAA supply to decapitated plants of rose consisted in placing a 2 ml-tube containing a basic medium (1% agar, 2.5 ml l^-1^ PPM, 100 mM mannitol), supplemented with NAA, at the cut end of the decapitated stem. Sucrose and mannitol were supplied through the rachis as described in (Lin *et al.*, 2011). The rachis was rapidly immersed in a sugar-containing liquid solution in a 1.5 ml reservoir. After 1 week the rachis was cut 0.5 cm lower. GR24 was supplied in aqueous solution (1.5% acetone) using the cotton-wick method described in (Corot *et al.*, 2017).

**Supplemental text 3. Estimation of the time at which growth starts.**

‘The time at which growth starts’ is defined as the time the bud starts rapidly elongating. It was estimated, for each individual bud, from the daily time course of bud length logarithm, in two steps. In a first step, one linear function was fitted for the phase of slow growth and one for the phase of rapid elongation using the lm procedure of R software for Windows. In a second step, the time when the two linear functions intersect was calculated, giving the time the bud starts rapidly elongating.

**Supplemental text 4. Sugar content determination**.

Internodes were harvested 24h after plants had been decapitated. Individual internodes were identified by their rank from shoot top. They were frozen in liquid nitrogen, lyophilized, and ground to a fine powder. Seventeen mg powder were homogenized with 1.3 ml of 80% aqueous ethanol at 80°C for 30 min and then, with 700 µl of 50% aqueous ethanol at 80°C for 30 min. The resulting suspension was centrifuged at 5500 rpm for 5 min at 4°C. The supernatant was used for soluble sugar analysis and the pellet for starch analysis. The supernatant was collected and concentrated using a Speed-Vac concentrator, until total ethanol removal, and then diluted in water to a final volume of 600 µl. This volume was used for sucrose, D-glucose and D-fructose content determination using a Konelab 20i sequential automat (Thermo Electron, Vantaa, Finland) and the ENZYTEC^TM^ system (Diagnostics, Viernheim, Germany).

The pellet was heated for one night to 60°C and then resuspended with 0.2 ml of 80% aqueous ethanol. The starch in the sample was then hydrolysed as follows. The sample was gelatinized by heat treatment at 100°C for 6 min in the presence of 2.9 ml of MOPS (Sigma Aldrich) buffer and 0.1 ml of thermostable α-amylase (3000 U.ml^-1^, Megazyme International Ireland). After cooling to 50°C, the sample was hydrolyzed for 30 min with 4 ml of sodium acetate (200 mM; pH 4.5) and 0.2 ml of amyloglucosidase (200 U.ml^-1^, Mégazyme International Ireland). 2 ml of supernatant were centrifuged at 10000 rpm for 10 min at 4°C. 600 µl were used for D-glucose content determination as described above.

**Supplemental tables**

**Supplemented table 1. Growth environment for each experiment.**

| ***In vitro* experiments** | | |  |
| --- | --- | --- | --- |
|  | Initial plants* | |  |
|  |  | Rose | Temperature-controlled greenhouse |
|  |  | Pea  Fig. **2e, 3c, 4f** | Growth chamber  130 μmol m^-2^ s^-1^; light/dark 16/8h photoperiod; 22/20°C at day/night |
|  |  | Pea  Fig. **4b** | Growth chamber  150 μmol m^-2^ s^-1^; light/dark 16/8h photoperiod; 22°C |
|  | Nodal segments | |  |
|  |  | Rose | Growth chamber  120 μmol m^-2^ s^-1^; a light/dark 16/8h photoperiod; 22/20°C at day/night |
|  |  | Pea WT  Fig. **2e, 3c, 4f** |  |
|  |  | Pea WT  Fig. **4b** | Growth chamber  150 μmol m^-2^ s^-1^; light/dark 16/8h photoperiod; 22°C |
|  |  | Pea *rms3*  Fig. **4d** | Growth chamber  70-80 μmol m^-2^ s^-1^; a light/dark 16/8h photoperiod; 22/20°C at day/night |
| **Decapitated rose plants** | | |  |
| Fig. **2a, 4c** | | | Growth chamber;  350 μmol m^-2^ s^-1^ at the top of the plants; light/dark 16/8h photoperiod; 22/19°C and 26/20°C at day/night for Fig. **2c** and Fig. **4c** respectively.  Water and mineral nutrition were provided each day by sub-irrigation. |
| Fig. **2b** | | | Growth chamber;  420 μmol m^-2^ s^-1^ at the top of the plants; light/dark 16/8h photoperiod; 22/20°C.  Water and mineral nutrition were provided each day by sub-irrigation. |

* plants from which nodal segments had been excised

**Supplemental table 2. Cytokinin quantification: retention times, limit of quantification (LOQ), and limit of detection (LOD)**

|  | Fig. **3a** | | | Fig. S2 | | |
| --- | --- | --- | --- | --- | --- | --- |
| Hormone | Retention time (min) | LOQ  (ng g^-1^ DW) | LOD  (ng g^-1^ DW) | Retention time (min) | LOQ  (ng g^-1^ DW) | LOD  (ng g^-1^ DW) |
| ZRMP  ^2^H_5_-t-ZRMP | 3.42  3.39 | 9.11 | 2.73 | 2.84  2.81 | 2.74 | 0.82 |
| ZR  ^2^H_5_-t-ZR | 4  3.97 | 0.67 | 0.2 | 3.84  3.82 | 0.42 | 0.13 |
| iPRMP  ^2^H_6_-iPRMP | 5.8  5.75 | 2.16 | 0.07 | 4.79  4.74 | 1.37 | 0.41 |
| iPR  iPR^•^ | 6.1  6.1 | 1.16 | 0.35 | 5.93  5.93 | 1.29 | 0.39 |
| iP  ^15^N-iP | 5.08  5.03 | 0.39 | 0.12 | 5.03  4.98 | 0.24 | 0.07 |

**Supplemented table 3. Definition, units, and estimated values of the model parameters.**

|  | **Parameter** | **Unit** | **Definition** | **Values** |
| --- | --- | --- | --- | --- |
| ***CK*** |  |  |  |  |
|  | *c*_1_ | mol.s^-1^ | Synthesis rate of *CK* without sucrose and auxin | 0.79 |
|  | *b*_1_ | mol^-1^ | Strength of *CK* synthesis inhibition by auxin | 0.96 |
|  | *a*_1_ | mol.s^-1^ | Maximum induction of *CK* synthesis by sucrose | 0.25 |
|  | *k*_1_ | mol^2^ | Parameter of the Hill function relating sucrose and *CK* synthesis | 0.19 |
|  | *d*_1_ | s^-1^ | *CK* degradation rate | 0.99 |
| ***SL*** |  |  |  |  |
|  | *c*_2_ | mol.s^-1^ | Base synthesis rate of *SL* without auxin | 0.34 |
|  | *a*_2_ | mol.s^-1^ | Maximum induction of *SL* synthesis by auxin | 24.89 |
|  | *k*_2_ | mol^2^ | Parameter of the Hill function relating auxin and *SL* synthesis | 294.58 |
|  | *d*_2_ | s^-1^ | *SL* degradation rate | 0.86 |
| ***I*** |  |  |  |  |
|  | *c*_3_ | mol.s^-1^ | Base production rate of *I* | 0.33 |
|  | *a*_3_ | mol^-1^.s^-1^ | Parameter relating the production rate of *I* to *SL* and sucrose | 5.64 |
|  | *u*_1_ | mol^-2^ | Minimum inhibiting effect of sucrose on *SL* response | 4.8 x 10^-13^ |
|  | *u*_2_ | mol^-4^ | Strength of sucrose inhibiting effect on *SL* response | 7.10 |
|  | *a*_4_ | mol.s^-1^ | Parameter relating the production rate of *I* to *CK* | 287.53 |
|  | *k*_3_ | mol^-2^ | Strength of *CK* effect on *I* production | 1000 |
|  | *d*_3_ | s^-1^ | Constant degradation rate of *I* | 0.99 |
| ***T*** |  |  |  |  |
|  | *m*_0_ | day | Intercept of the linear relationship between *T* and *I* | -2.2 |
|  | *m*_1_ | day. mol^-1^ | Sensitivity of the time at which elongation starts to *I* | 3.5 |
|  | *I*_0_ | mol | Threshold of *I* above which bud elongation is completely prevented | 3 |

**Supplemental table 4. Values of the variables used for model calibration.**

| Model  variables | Values | | | | | | | | | | | | | | | |
| --- | --- | --- | --- | --- | --- | --- | --- | --- | --- | --- | --- | --- | --- | --- | --- | --- |
|  | 0 μM NAA | | | | 1 μM NAA | | | | | | 2.5 μM NAA | | | | | |
|  | Sucrose (mM) | | | | Sucrose (mM) | | | | | | Sucrose (mM) | | | | | |
|  | 10 | 50 | 100 | 250 | 10 | 50 | | 100 | | 250 | 10 | 50 | | 100 | | 250 |
|  |  |  |  |  |  | -SLs | +SLs | -SLs | +SLs |  |  | -CKs | +CKs | -CKs | +CKs |  |
| *S* | 0.1 | 0.5 | 1 | 2.5 | 0.1 | 0.5 |  | 1 |  | 2.5 | 0.1 | 0.5 |  | 1 |  | 2.5 |
| *A* | 0 | 0 | 0 | 0 | 1 | 1 |  | 1 |  | 1 | 2.5 | 2.5 |  | 2.5 |  | 2.5 |
| *SL* | 0.4 | 0.4 | 0.4 | 0.4 | X** | X** | X**  +10 | X** | X**  +10 |  | 1 | 1 |  | 1 |  | 1 |
| *CK*^a^ | 0.75 | 0.97 | 1.00 | 1.10 | 0.51 | 0.59* |  | 0.58* |  |  | 0.23* | 0.36* | 0.61 | 0.46 | 0.71 | 0.51 |
| *T*^b^ | 3.2 | 2.7 | 1 | 0.9 |  | 5.8 |  | 3.5 |  | 2.1 |  |  |  |  |  | 3.1 |

^a^Measured values of iP content.

^b^Calculated from bud elongation kinetics as described in supplemental text 3.

*Measured relative values of iP concentration, presented Fig. S2, multiplied by 1.35, a scaling factor calculated so that iP concentration measured with 2.5 μM NAA and 100 mM sucrose observed Fig. S2 matches with that observed Fig. **3a**. Values are represented relative to the treatment 100 mM sucrose and 0 μM NAA.

**X is estimated after parameter optimization.

**Supplemental figures**

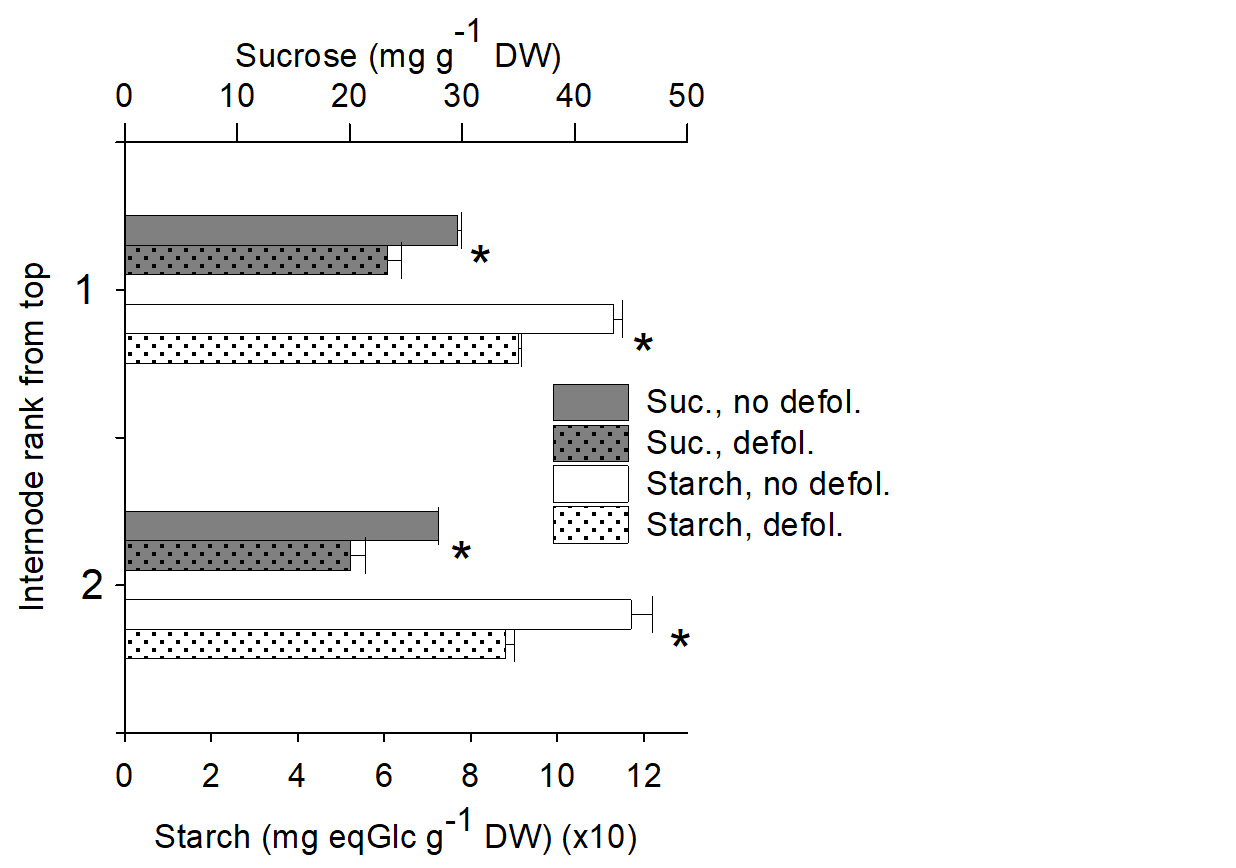

**Fig. S1. The level of defoliation of decapitated plants modulates sugar level.** Sucrose (Suc.) and starch content of the two topmost internodes of decapitated plants of rose supplied at their top with 0 or 10 μM NAA, in case of non-defoliation (no defol.) or partial defoliation (defol.). Data are means ± SEM (*n*=3 pools of 2 plants). Asterisks indicate significant differences between defoliation treatments (Student’s test; *p*<0.05). The method used for sugar quantification is described in supplemental text 4.

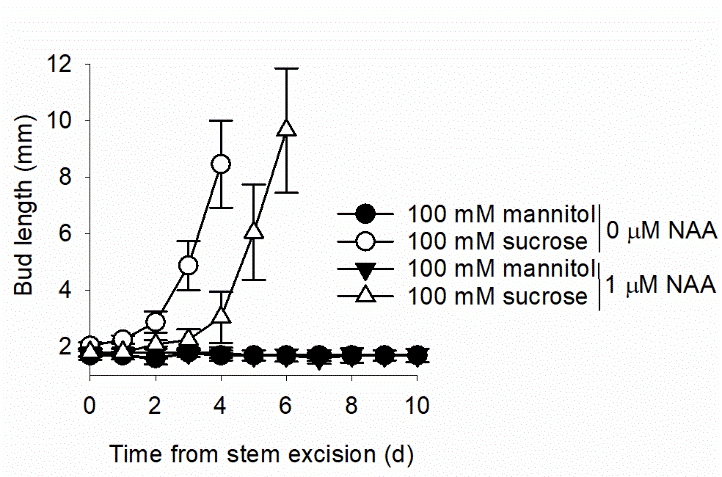

**Fig. S2. Sucrose antagonizes the inhibiting effect of auxin on bud outgrowth for rose nodal stems *in vitro*.** Bud elongation in response to 100 mM sucrose or 100 mM mannitol, an osmotic control, without or with 1 μM NAA. NAA, sucrose, and mannitol were supplied in the growth medium of isolated nodal stems of rose. Data represent the bud with the median final length (*n*=10). Error bars represent 95% confidence intervals.

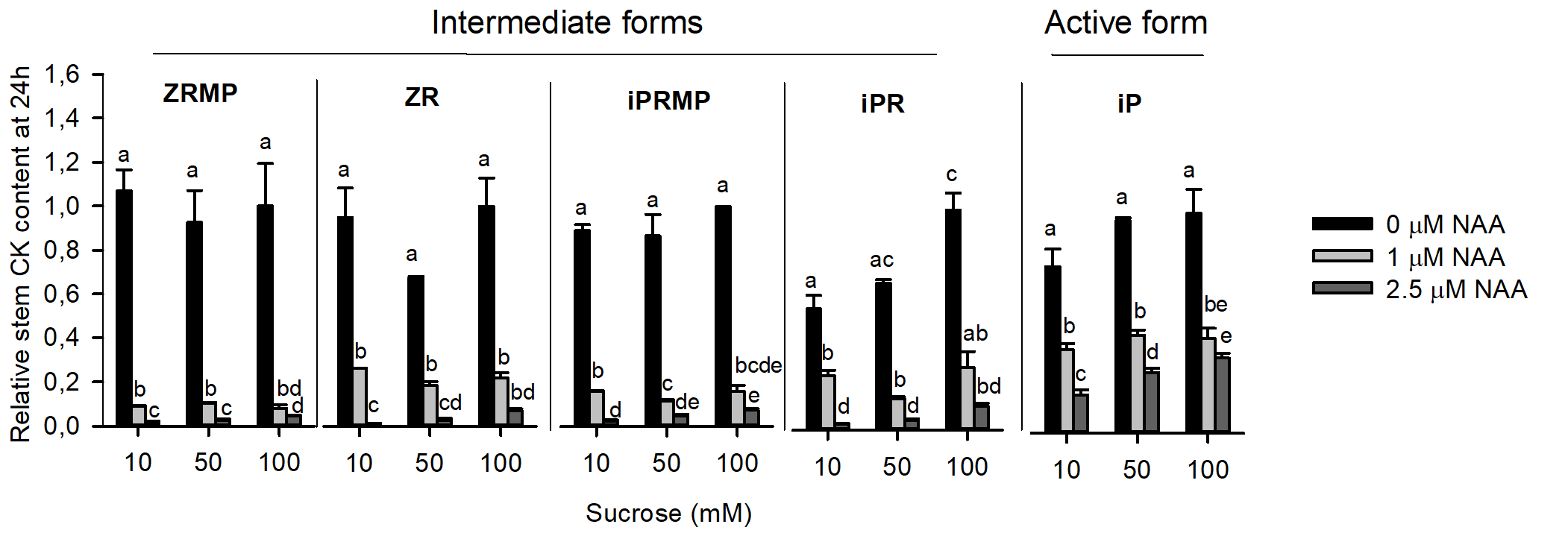

**Fig. S3. Cytokinin levels of the nodal stem strongly change with auxin, but not with sucrose.** Response of nodal cytokinins (CK) to different levels of NAA and sucrose for nodal segments of rose grown *in vitro*. The different cytokinin (CK) forms were quantified 24h after nodal stem excision (active Z form not detected). Data are means ± SEM (*n*=3 pools of 3 stem segments). Values are represented relative to the treatment 100 mM sucrose and 0 μM NAA. Different letters indicate significant difference between means (Student’s test; *p*<0.05). The protocol used for cytokinin quantification is similar to that used for Fig. **3a**, except that 0.5 ng of each internal standard was used and the dry extracts were dissolved in 100 µl (instead of 140 µl).

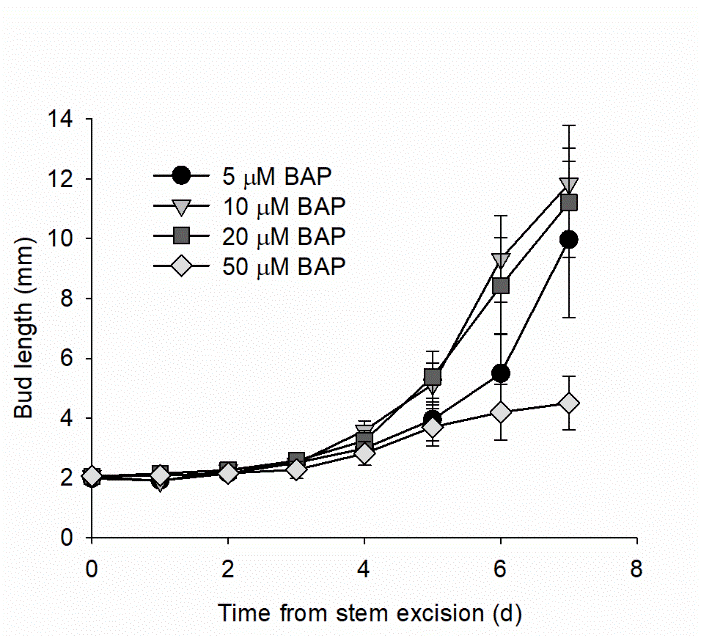

**Fig. S4. 10 μM BAP optimally stimulates bud outgrowth for rose nodal segments *in vitro*.** Effect of different levels of BAP in the medium on bud elongation kinetics for nodal segments of rose grown *in vitro* with 100 mM sucrose and 2.5 μM NAA. Data represent the bud with the median final length (*n*=10). Error bars represent 95% confidence intervals.

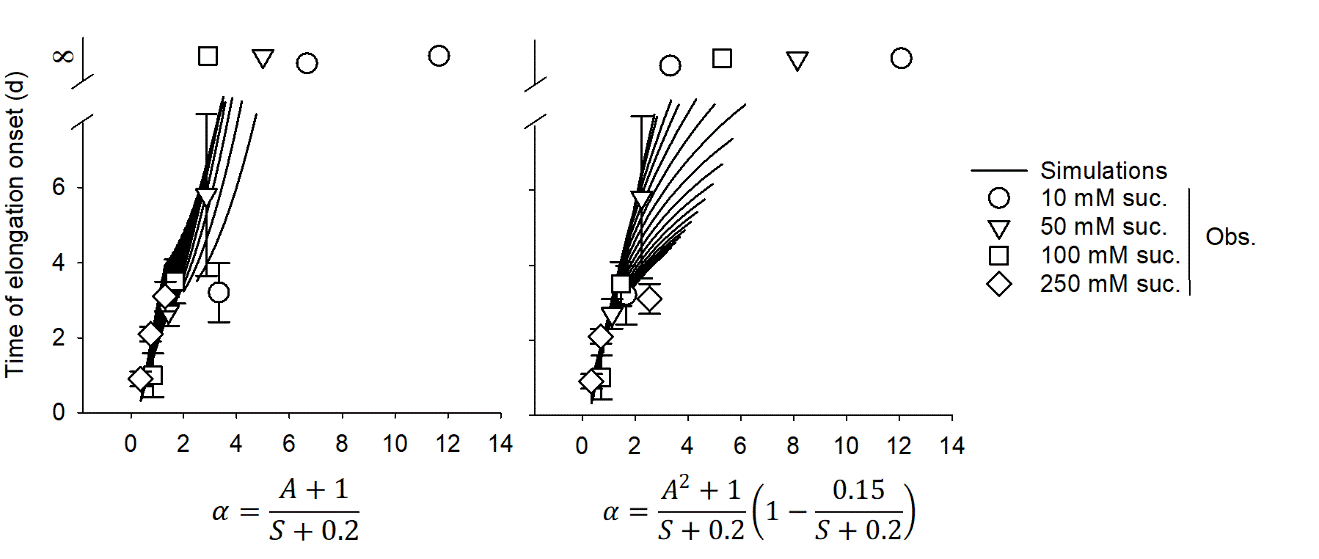

**Fig. S5. Deviations from the one-to-one relationship after changes of the control variable *α*.** Relationship between (i) the time at which elongation starts and (ii) two different formulae of *α*. In the first formula (left), *α* is the ratio between affine functions of auxin and sucrose levels ($\frac{A+1}{S+0.2}$). In the second formula, *α* is the ratio between affine functions of square auxin level and sucrose level ($\frac{A^{2}+1}{S+0.2}$), modulated by a sucrose-dependent correcting term $\left( 1-\frac{0.15}{S+0.2} \right)$. The different lines show simulations for different sucrose levels and the different symbols the biological observations for nodal segments of rose *in vitro* grown at different sucrose levels (Obs.). Observations of an absence of bud elongation are represented by a time at which elongation starts which is infinite (∞). For simulations, bud elongation is completely prevented above a threshold of 8.3 days.

**Brewer PB, Dun EA, Gui RY, Mason MG, Beveridge CA. 2015.** Strigolactone Inhibition of Branching Independent of Polar Auxin Transport. *Plant Physiology* **168**(4): 1820-U1215.

**Corot A, Roman H, Douillet O, Autret H, Perez-Garcia MD, Citerne S, Bertheloot J, Sakr S, Leduc N, Demotes-Mainard S. 2017.** Cytokinins and Abscisic Acid Act Antagonistically in the Regulation of the Bud Outgrowth Pattern by Light Intensity. *Frontiers in Plant Science* **8**.

**Lin YH, Lin MH, Gresshoff PM, Ferguson BJ. 2011.** An efficient petiole-feeding bioassay for introducing aqueous solutions into dicotyledonous plants. *Nature Protocols* **6**(1): 36-45.
